# Short-term auditory priming in freely-moving mice

**DOI:** 10.1101/2023.04.03.535329

**Authors:** Shir Sivroni, Hadas Sloin, Eran Stark

## Abstract

Priming, a change in the mental processing of a stimulus as a result of prior encounter with a related stimulus, has been observed repeatedly and studied extensively. Yet currently there is no behavioral model of short-term priming in lab animals, limiting the study of the neurobiological basis of priming. Here, we describe an auditory discrimination paradigm for studying response priming in freely-moving mice. We find a priming effect in success rate in all mice tested on the task. In contrast to humans, we do not find a priming effect in response times. Compared to non-primed discrimination trials, the addition of incongruent prime stimuli reduces success rate more than congruent prime stimuli, suggesting a cognitive mechanism based on differential interference. The results establish the short-term priming phenomenon in rodents, and the paradigm opens the door to studying the cellular-network basis of priming.

## Introduction

We are all familiar with the feeling that related but irrelevant experiences unintentionally influence our decisions. The way products, ideas, and behaviors are promoted or presented affects our tendencies or desires^1–3^. For example, if you own a restaurant and have ordered too much French wine, an efficient approach for selling that specific wine may involve playing French music in the background^4^. The mental process behind the effect is called “priming” ^1,2,5^. Priming has been studied in many fields including political science^6^, cognitive science^7–10^, economics^2,11,12^, psychophysiology^13^, developmental psychology^14–16^, psycholinguistics^17,18^, law^19^, education^20,21^, philosophy of the mind^22,23^, and business^24^. When the effect is prolonged priming relies on long-term memory^25^, whereas if the effect is short-lived (e.g., seconds), priming relies on short-term-memory^26^. Here, we focus on short-term priming.

Formally, priming is defined as a change in the mental processing of a stimulus as a result of prior encounter with a related stimulus^27,28^. In the restaurant example, the type of music played would be the “prime” experience, and the type of wine selected would be the “target”. Multiple paradigms have been developed for investigating short-term priming^28–31^. A distinction is made when the prime is processed consciously, i.e., the subject is aware of the prime stimulus, as opposed to unconsciously^28,32–34^. One specific paradigm is “response priming”^35,36^, in which the prime and target stimuli are presented in quick succession and are coupled with identical or alternative responses. Response priming studies typically employ two alternative forced choice (2AFC) tasks for quantifying the influence of the prime on performance. Even when the subject is told that the prime is irrelevant for the performance of the task, the prime influences the outcome^28^ (**Fig. 1A**). For example, the word “chair” (here, the target stimulus) is classified as “artificial” more rapidly when following the word “table” (the congruent prime stimulus; **Fig. 1A, left**), as opposed to following the word “sky” (the incongruent prime stimulus; **Fig. 1A, right**). The priming effect is quantified by a performance difference between congruent prime trials (CPT) and incongruent prime trials (IPT). Differences are typically evident in response times^27,32,37^ and may also be evident in success rates^36,38^.

**Figure 1.**
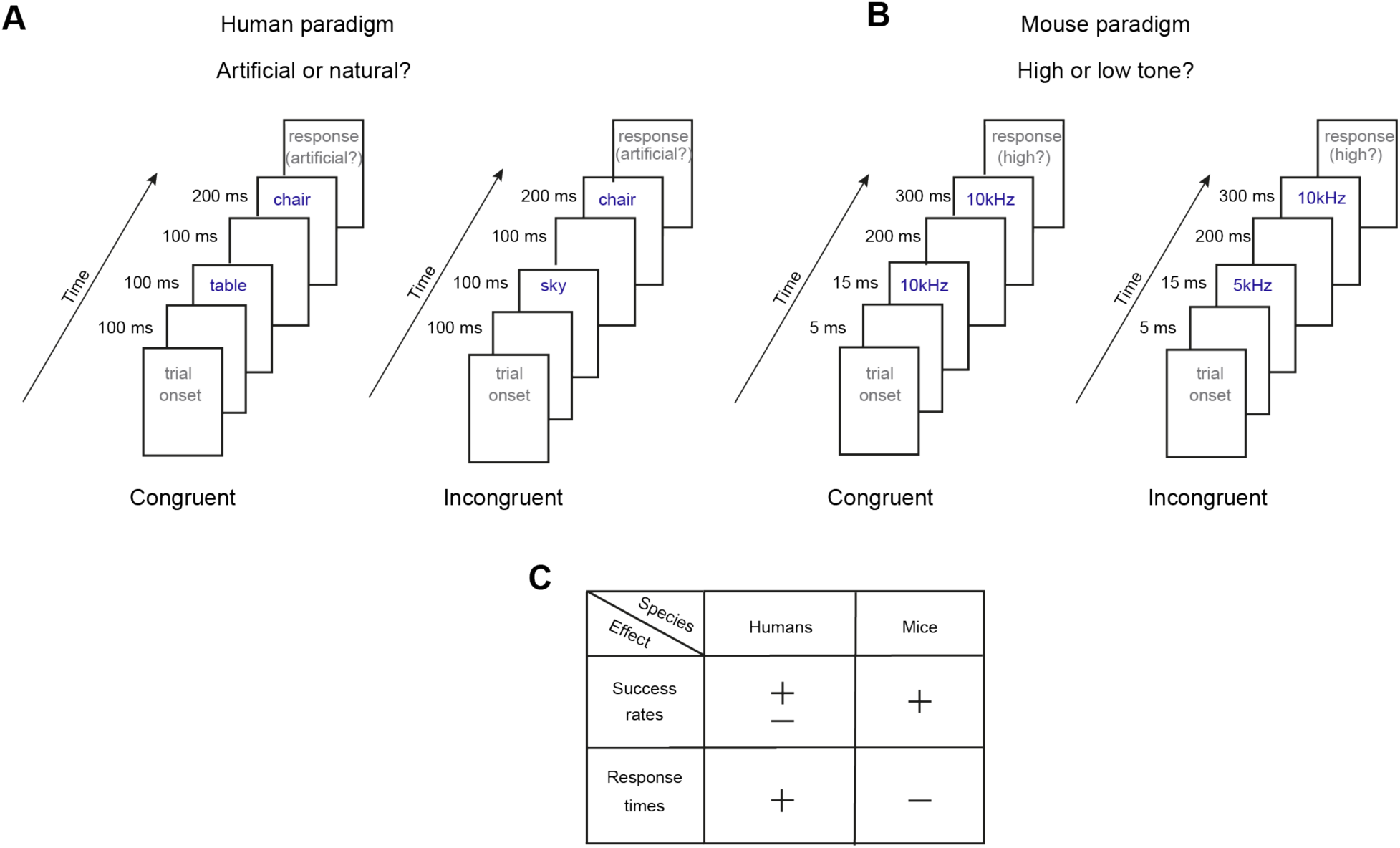
Paradigms for studying short-term priming in humans and in mice. **(A)** Standard paradigm for testing short-term response priming in humans. **Left**, A typical timeline for a congruent prime trial (CPT). “Table” and “chair” are both pieces of furniture and belong to the same semantic content group. **Right**, A typical timeline for an incongruent prime trial (IPT). In the context of the experiment, the word “sky” (the prime stimulus) and the word “chair” (the target stimulus) do not belong to the same semantic content group. A “sky” target stimulus implies a “left” choice, and “chair” implies “right”. When the prime and target stimuli are congruent, human response times are typically shorter. **(B)** Proposed paradigm for testing short-term priming in mice. **Left**, A typical timeline for a CPT. The prime stimulus (short pure tone) and the target stimulus (long pure tone) belong to the same frequency group, since both tones have the same frequency, 10 kHz. **Right**, A typical timeline for an IPT. In the context of the experiment, 5 kHz and 10 kHz pure tones do not belong to the same content group. A target stimulus of “5 kHz” implies a “left” choice, and “10 kHz” implies a “right” choice. When the prime and target stimuli are congruent, mouse success rates are typically higher. **(C)** In humans, the typical priming effect is in response time. In contrast, we find that in mice the priming effect is in success rates, and not in response times.

Almost all research on priming has been conducted on human subjects. With human subjects, verbal explanations are possible, but not with animals. An extensive literature review revealed less than a dozen studies related to short-term priming in animals: birds^39–43^, rats^44,45^, lemurs^46^, and monkeys^47,48^. To date, there are no studies of priming involving cellular-level electrophysiology in humans or animals. Consequently, existing models of priming are phenomenological, including “global neuronal workspace”^49^ and “accumulator”^50^ models. There are no physiological mechanistic models for priming^28,51^. For instance, priming may consist of a combination of behavioral repetition suppression and top-down influence of attentional amplification of task-congruent processing pathways^52^. However, alternative mechanistic models are difficult to contest without access to cellular-network activity.

Here, we developed an auditory response priming paradigm for freely-moving mice (**Fig. 1B**). In two preliminary stages, we train the subjects to respond to long pure tones and to ignore short pure tones. We then couple short and long tones in the same trial. In contrast to studies of short-term priming in humans, we do not find any difference in response times. However, we find a consistent difference between success rates during CPTs and IPTs in all subjects. The results show that priming occurs in mice, suggesting that mice enjoy the same benefit as humans from the automatic activation of a first stimulus when performing a task on the second. Finally, we find that compared to same-session discrimination trials, success rates are reduced during IPTs more than during CPTs, suggesting that differential interferences underlies short-term priming in the mice. The presented model system opens the door to studying the molecular, genetic, and cellular-network mechanisms at the basis of priming.

## Results

All behavioral training and testing sessions were carried out in the same apparatus (**Fig. 2A**). Sessions were organized in three stages, each consisting of one or more distinct tasks (**Fig. 2B**). The first stage included discrimination sessions, composed of discrimination trials (DTs; **Fig. 2CD, top; video1**). The second stage consisted of validation sessions, in which DTs were supplemented by validation trials (VTs; **Fig. 2CD, middle; video2**). The third stage consisted of priming sessions, in which the DTs were supplemented by priming trials (PTs; **Fig. 2CD, bottom; video3**). To move from the first to the second stage, or from the second to the third stage, the animals had to fulfil a performance criterion. The third stage did not include training phases, and only involved testing on the priming task.

**Figure 2.**
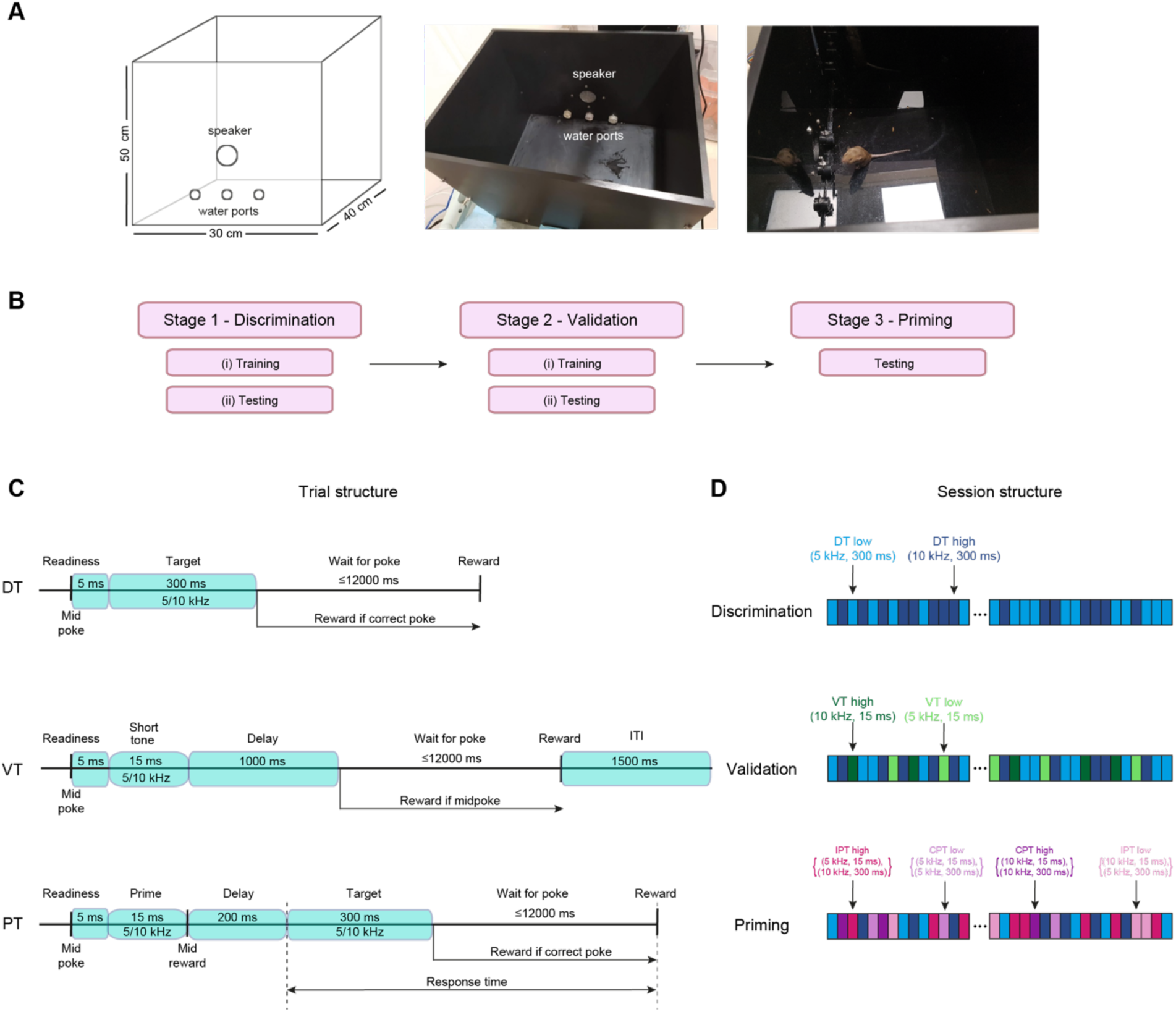
Apparatus and training procedure for short-term auditory priming in mice. **(A)** The behavioral apparatus. One wall of the box contains all sensors and actuators, including a speaker and three illuminated water ports equipped with infrared LEDs for detecting nose pokes. **(B)** Mice are trained in two preliminary stages before being tested on the priming task. Stage 1 includes a simple discrimination task, in which the stimulus-response (tone-side) contingency is learned using long (300 ms) pure tones. Stage 2 includes a validation task, in which the mice learn that short (15 ms) pure tones are irrelevant for choosing a correct response. Stage 3 includes the priming task. **(C)** Timelines for the three types of trials. **Top,** Discrimination trials (DTs). **Middle**, Validation trials (VTs). **Bottom**, Priming trials (PTs). ITI, inter-trial interval. **(D)** Example sequence of trials during discrimination, validation, and priming sessions. Every rectangle represents a single trial. See also **video1** (an excerpt from a discrimination session), **video2** (a validation session), and **video3** (a priming session).

### Mice learn to discriminate lower pure tones from higher pure tones

The first stage of the learning process contained only discrimination sessions^53^. In each DT, one of two pure tones was presented for 300 ms: a low target tone (5 kHz) or a high target tone (10 kHz). Both tones are in the audible range for mice^54^ and for humans. During the training period, the mouse had to associate the low frequency tone with the left port, and the high frequency tone with the right port. An example post-learning discrimination session can be found in **Fig. 3A**. The color represents the target stimulus frequency, which can be 5 kHz (light blue) or 10 kHz (dark blue). The height represents the response (mouse choice), which can be the left port (bottom), ignoring the tone (middle), or the right port (top). In the example session there were 154 trials with 5 kHz tones and 157 trials with 10 kHz tones. The mouse ignored two trials towards the end of the session, and was incorrect in 29 other trials, yielding an overall success rate of 280/311 (90%).

**Figure 3.**
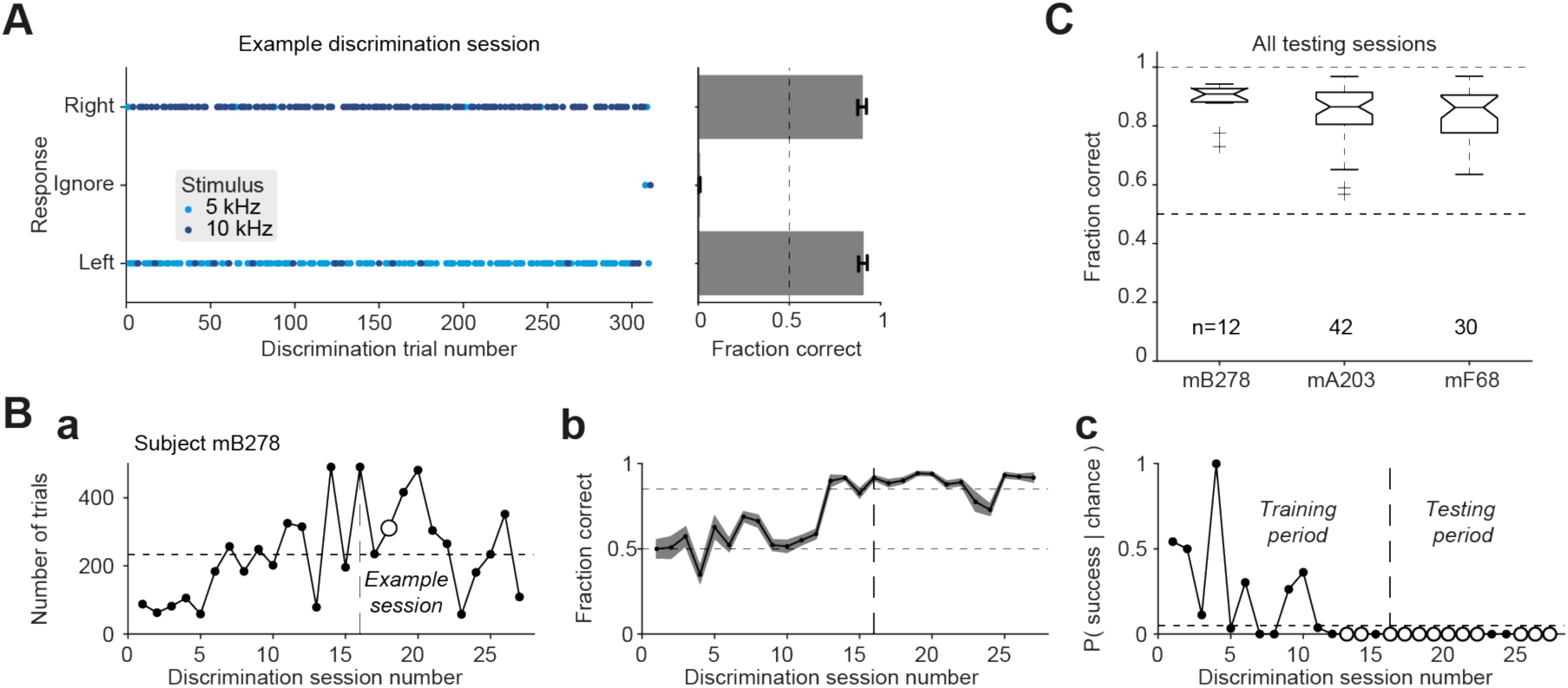
Mice learn to discriminate lower pure tones from higher pure tones. **(A)** An example discrimination session performed by subject mB278. **Left**, Every dot represents a single DT. Dot color represents the target stimulus in the trial, i.e., the frequency of the 300 ms tone, and the vertical location represents the response (choice) during the trial. Possible responses include the left port, ignorance, and the right port. A session with perfect performance would yield a graph with all dark blue dots on top, all light blue dots at the bottom, and no dots in the middle. The example session consisted of 311 trials with a 90% success rate. The mouse ignored one low and one high tone, both towards the end of the session. **Right**, Fraction histograms for left correct trials (bottom), right correct trials (top), and ignored trials (middle). **(B)** All discrimination sessions carried out by one subject (mB278). The vertical dashed line marks the first of four consecutive successful sessions, considered the point at which the mouse has learned the discrimination task. The same session is the first session of the testing period (panel **C**). **a** The number of trials in every discrimination session. **b** Success rate, defined as the fraction of correct DTs. **c** The p-values of deviation from chance for every session (Binomial test comparing to chance level, 0.5). Successful sessions (p<0.05 and success rate above 0.85) appear as enlarged empty circles. **(C)** Post-learning performance of subjects mB278, mA203, and mF68 on the discrimination task. Every box plot shows median and interquartile range (IQR) for one subject. Whiskers extend for 1.5 times the IQR in every direction, and a plus sign indicates an outlier. All mice learned the task, with an overall success rate of 88% [79% 91%] (median [IQR], n=84 sessions).

Mice started the first discrimination sessions with success rates around 0.5. During training, the number of trials per session (**Fig. 3Ba**) and the success rates (**Fig. 3Bb**) gradually increased. All mice tested on the paradigm learned the discrimination task, achieving the criterion of at least four consecutive sessions with above chance performance (p<0.05, Binomial test) and success rate above 85%. For the example subject, the transition from learning to post-learning is marked by a dashed vertical line in **Fig. 3B**. The number of training sessions before learning were 15, 13, and 24, for subjects mB278, mA203, and mF68, respectively. After the learning period ended, additional discrimination sessions were conducted to measure and consolidate performance. During the post-learning sessions, all subjects exhibited consistently high performance (**Fig. 3C**), with a median [interquartile-interval, IQR] post-learning success rate of 88% [79% 91%] (n=84 sessions). The number of trials was self-paced by the mice, with a median [IQR] of 239 [171 322] trials per post-learning discrimination session. Subject-specific success rates were 91% [88% 92%] (n=12 sessions, mB278); 87% [80% 91%] (n=42, mA203); and 86% [76% 90%] (n=30, mF68). Thus, all the mice achieved stable and high performance of discriminating between lower (5 kHz) and higher (10 kHz) pure tones.

### Mice learn to ignore short tones while discriminating long tones

After learning the discrimination task and exhibiting consistently-high performance, the mice were trained to ignore short (15 ms) pure tones. A crucial element of every testable priming paradigm is that the subject will understand that the task should be performed on the target and not on the prime. With human subjects, verbal explanations are useful, but not with mice. To establish that short tones do not predict a correct choice of lateral port (right or left) and are therefore irrelevant, validation trials (VTs) were used. In addition to VTs, every validation session contained DTs, identical to the DTs employed during discrimination sessions. In each VT, one of two pure tones was presented for 15 ms: a low short tone (5 kHz), or a high short tone (10 kHz). During training, the mouse had to learn that there is no association between short tones and a choice of a side port. In practice, the subject had to respond to short tones at chance level, or ignore the short tones and choose a side port only after long tones.

All the VTs of an example post-learning validation session are shown in **Fig. 4A**. The colors represent the stimulus (short tone frequency, 5 or 10 kHz), and the height represents the response (left port, ignorance, or right port). The example session included a total of 353 trials, of which 82 were VTs and the rest were DTs. Of the VTs, 38 trials included low short tones (5 kHz; light green) and 44 trials included high short tones (10 kHz; dark green). The mouse correctly ignored 35/38 (92%) of the low tones and 36/44 (82%) of the high tones. Thus, 71/82 (87%) of the VTs were ignored. During not-ignored trials, the “success rate” was 10/11 (91%). To take into account the fraction of the ignored trials as well as the responses during not-ignored trials, we defined an “irrelevance score” (**Fig. 4B**). The irrelevance score is close to 1 when the short tones are ignored (**Fig. 4B, red**) and close to 0 when the mouse fails to ignore the short tones (**Fig. 4B, blue**). For the example validation session, the irrelevance score was 0.89.

**Figure 4.**
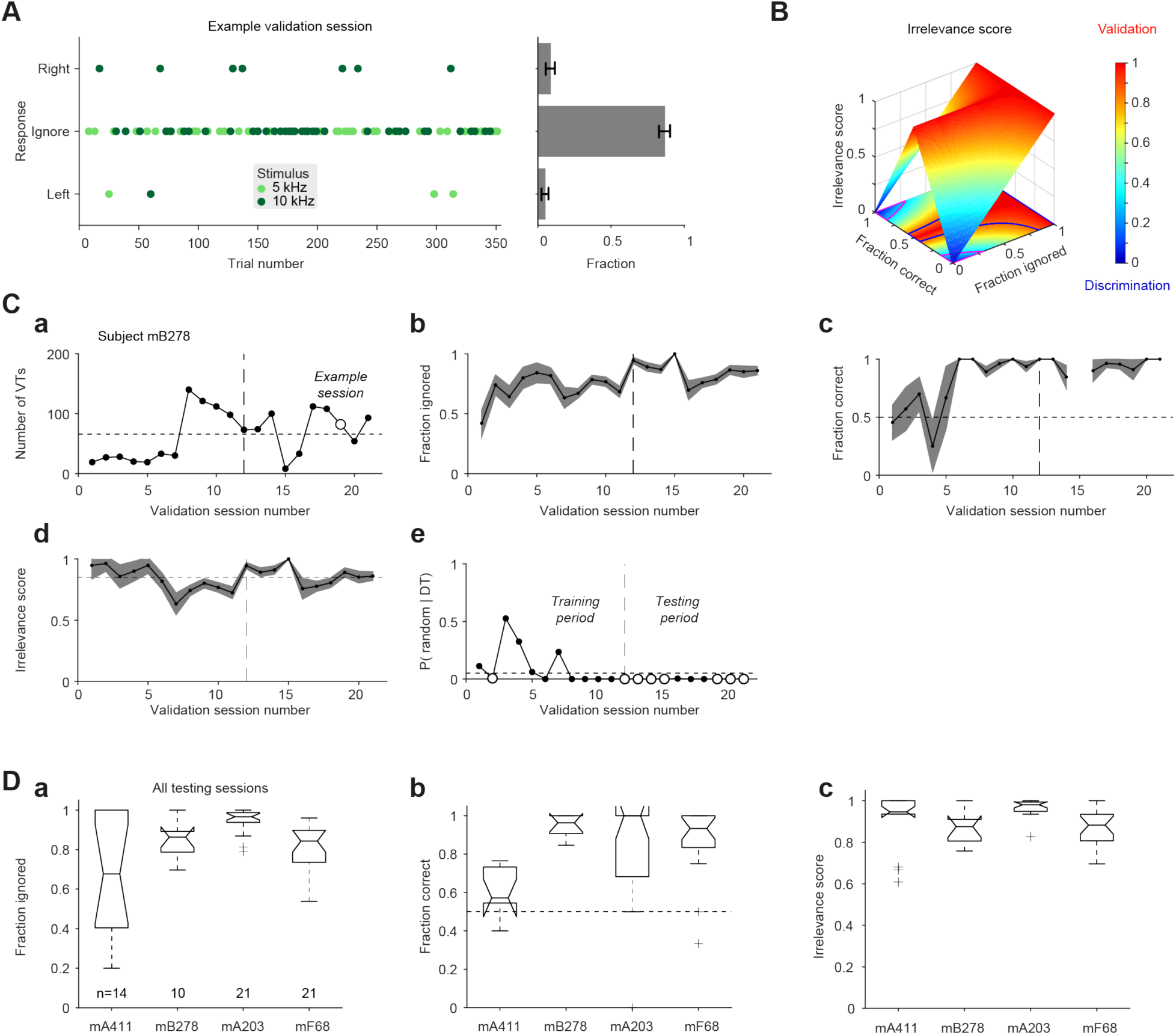
Mice learn to ignore short tones while discriminating long tones. **(A)** All VTs in an example validation session of subject mB278. **Left**, Every dot represents a single VT. Dot color represents the frequency of the short pure tone (15 ms), and the vertical location of the dot represents the response. A validation session with perfect performance during VTs should yield a graph with all dots in the middle. **Right**, Fraction histograms for the number of the trials corresponding to every response. The subject ignored short tones during 71/82 (87%) of the VTs. Of the not-ignored VTs, the mouse made a “correct” choice in all three low short tones and in 7/8 of the high short tones. **(B)** Dependence of the “irrelevance score” on the fraction of ignored VTs and on success during not-ignored VTs. When performance is very close to discrimination, scores fall in blue-hued parts of the graph, indicating that the subject has not learned the validation task. When the desired performance on the validation task is achieved, scores fall in the red-hued parts. **(C)** Performance of subject mB278 during all validation sessions. The vertical dashed line marks the first of four consecutive successful sessions, considered the point at which the mouse learned the validation task. **a** The number of VTs in every validation session. **b** The fraction of ignored VTs. **c** The fraction of “correct” trials out of the not-ignored VTs. **d** The irrelevance score. **e** The p-values of irrelevance score for every validation session (Monte Carlo test). **(D)** Post-learning performance on the validation task, pooled over all testing sessions for subjects mA411, mB278, mA203, and mF68. Box plot conventions are the same as in Fig. 3C. **a** Fraction of ignored VTs pooled over all post-learning sessions for every mouse. **b** “Success rates” for the not-ignored VTs. **c** Irrelevance scores. All mice learned the validation task, with an overall irrelevance score of 0.94 [0.86 0.99] (median [IQR], n=66 sessions).

The mice started training on the validation task after having learned the discrimination task to criterion. Thus, during the initial validation sessions the mice did not ignore the short tone stimuli (**Fig. 4Cb**). However, during training, the fraction of ignored VTs gradually increased and stabilized (**Fig. 4Cb**). Learning was defined as crossing a criterion of four consecutive sessions with above chance irrelevance score (p<0.05, Monte Carlo test) maintained above 0.85. For the example subject, the transition from learning to post-learning is marked by a vertical dashed line (**Fig. 4Cd**). During training, subjects mA411, mB278, mA203, and mF68, required 13, 11, 8, and 16 validation sessions. After the training period ended, additional validation sessions were conducted to consolidate and measure performance on the validation task (**Fig. 4D**). During the post-learning period, the mice consistently ignored the short tones (**Fig. 4Da**). The median [IQR] number of trials was 261 [173 366] per session, of which 52 [30 79] were VTs (n=66 sessions in four mice). Subject-specific irrelevance scores were 0.95 [0.68 1.00] (n=14 sessions, mA411); 0.88 [0.78 0.91] (n=10, mB278); 0.98 [0.95 0.99] (n=21, mA203); and 0.88 [0.77 0.93] (n=21, mF68; **Fig. 4Dc**). Overall, the post-learning irrelevance scores were 0.94 [0.86 0.99] (n=66). Thus, all mice learned to ignore the short tone stimuli.

### All mice show a priming effect in success rates

The third stage contained priming sessions. Every priming session consisted of DTs, identical to the DTs employed during discrimination sessions, and priming trials (PTs). Every PT included a short tone (the prime stimulus) followed by a long tone (the target stimulus; **Fig. 2CD, bottom**). PTs in which the short and long tones are of the same frequency are called congruent PTs (CPTs), whereas PTs in which the frequencies of the short and long tones differ are called incongruent PTs (IPTs). During PTs, the mouse was rewarded in the same manner as during DTs, namely for poking the right port following a low long tone, and for poking the left port following a high long tone.

All PTs of an example priming session are shown in **Fig. 5A**. The color represents the stimulus (long tone frequency, 5 or 10 kHz), and the height represents the response (left port, ignorance, or right port). The session included 234 trials, of which 91 were PTs and the rest were DTs. Of the PTs, 48 were CPTs (**Fig. 5A, left**) and 43 were IPTs (**Fig. 5A, right**). The mouse made a correct choice during 47/48 (98%) of the CPTs and during 35/43 (81%) of the IPTs. We quantified the “priming effect” in success rates by the difference between CPT and IPT success rates, which was 17% in the example session (p=0.011, permutation test). Thus, in the specific case there was a consistent effect of the short tones on choice to the long tones: a short-term priming effect.

**Figure 5.**
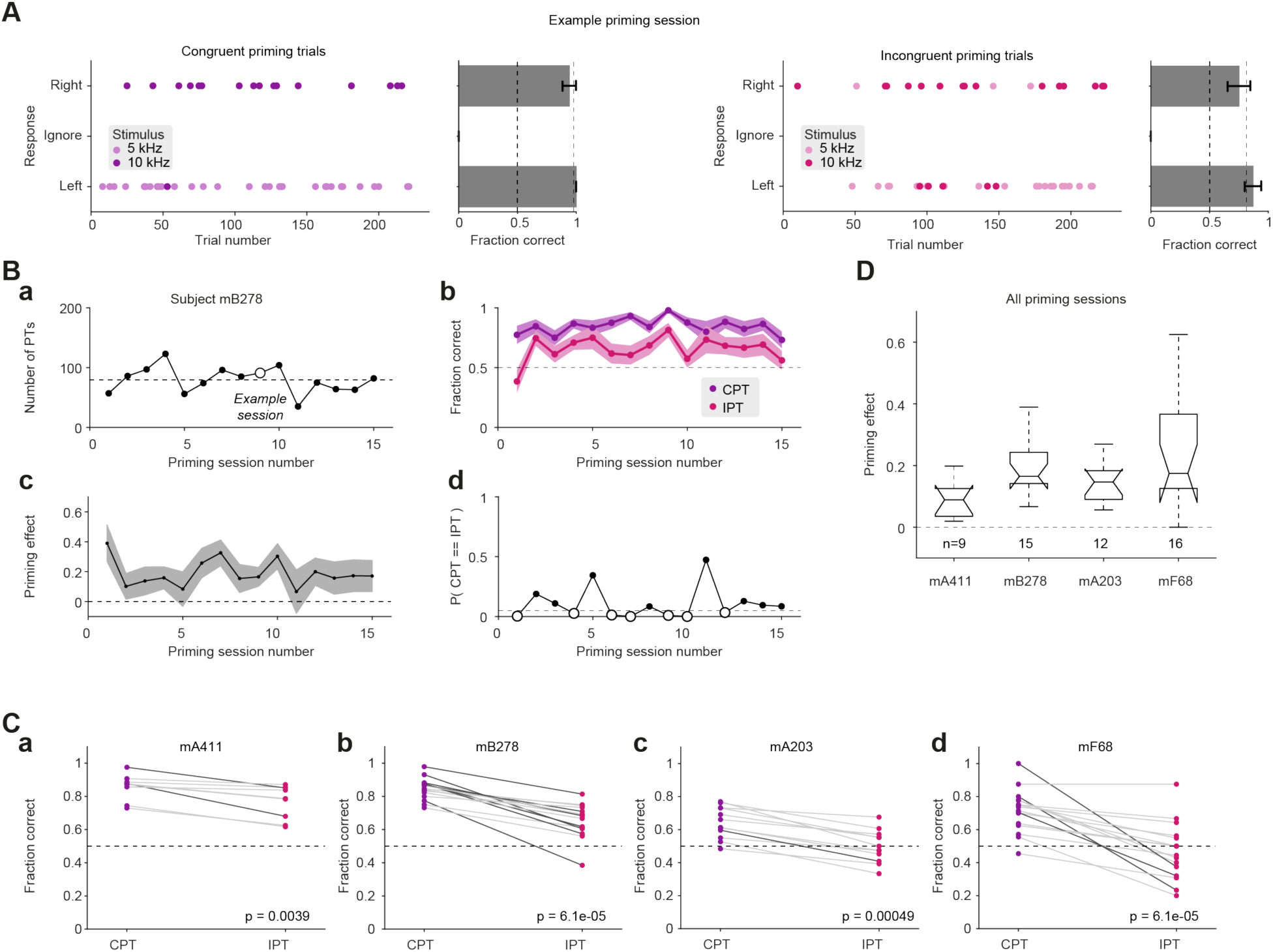
All mice show a priming effect in success rates. **(A)** All PTs during an example priming session performed by subject mB278. Every dot indicates a single PT. Dot color represents the frequency of the target stimulus (300 ms tone), and the vertical location of the dot represents the response (choice). A priming session with perfect performance would yield a graph with all dark-colored (violet or pink) dots on top and all light-colored (violet or pink) dots at the bottom. **Left**, All congruent PTs (CPTs) during the session. **Right**, All incongruent PTs (IPTs). Success rate is higher during CPTs (98%) compared with IPTs (81%; p=0.011, permutation test). **(B)** Performance during all priming sessions of subject mB278. **a** The number of PTs during every priming session. **b** Success rates during the CPTs (violet) and during the IPTs (pink) of every session. **c** The “priming effect” during every session, defined as the difference between the same-session CPT and IPT success rates. **d** The p-values of the priming effect for every priming session (permutation test, comparing to an effect size of zero). **(C)** Success rates for all priming sessions performed by subjects mA411, mB278, mA203, and mF68. Every violet dot represents the success rate during the CPTs during one session and is connected by a line with a pink dot, representing the success rate during the same-session IPTs. P-values, two-tailed Wilcoxon’s signed-rank paired test. **(D)** Priming effects for the four subjects. Box plot conventions are the same as in Fig. 3C. The overall priming effect is 15% [9.1% 20%] (median [IQR], n=52 sessions).

In contrast to the discrimination and validation tasks, we did not train the mice on any specific criterion during PTs. However, success rates were higher during CPTs compared with IPTs during all sessions of every mouse, with only one exceptional session (see below). For instance, subject mB278 exhibited a median [IQR] priming effect of 16% [10% 26%], with a range of [7%, 39%] (**Fig. 5Bbc**). The priming effect was significant only in some of the sessions (7/15 sessions with p<0.05, permutation test; **Fig. 5Bd**; dark grey lines in **Fig. 5Cb**). However, a positive priming effect was observed in all sessions of the same subject (15/15; p<0.001, Wilcoxon’s paired signed-rank test; **Fig. 5Cb**). Similar results were observed in the other mice: success rates were consistently higher during CPTs compared with IPTs (p<0.005 in all cases; **Fig. 5Ca,c,d**).

We conducted 52 priming sessions in four mice, with a median [IQR] of 148 [91 199] trials per session, of which 71 [34 91] were PTs. There was a significant priming effect in 14/52 (27%) of the sessions (dark grey lines in **Fig. 5C**). The effect was positive in 51/52 (98%) of the sessions. In one session, success rates were 88% during both CPTs and IPTs (**Fig. 5Cd**). Subject-specific priming effects were 8.9% [3.3% 12.5%] (n=9 sessions, mA411); 16.5% [10.1% 25.7%] (n=15, mB278); 14.7% [8.9% 17.9%] (n=12, mA203); and 17.4% [12.5% 35.6%] (n=16, mF68; **Fig. 5D**). Thus, all mice exhibited a consistent priming effect, with a global priming effect of 15.1% [9.1% 20%] (median [IQR] over n=52 sessions).

### None of the mice show a priming effect in response times

In human subjects, priming effects are typically observed in response times^28,55,56^ or in both response times and success rates^36,38,50^. To determine whether mice exhibit a priming effect in response times in addition to success rates, we defined response time as the duration from target tone onset to side port selection (**Fig. 2C**). For the same example session of subject mB278 described in **Fig. 5A**, the median [IQR] response times were 487 ms [414 623] ms during n=48 CPTs, and 522 ms [460 697] ms during n=43 IPTs (**Fig. 6A**). In the session, response times during CPTs and IPTs did not exhibit distinct medians (p=0.15, Mann-Whitney U-test) or distinct variances (p=0.49, permutation test).

**Figure 6.**
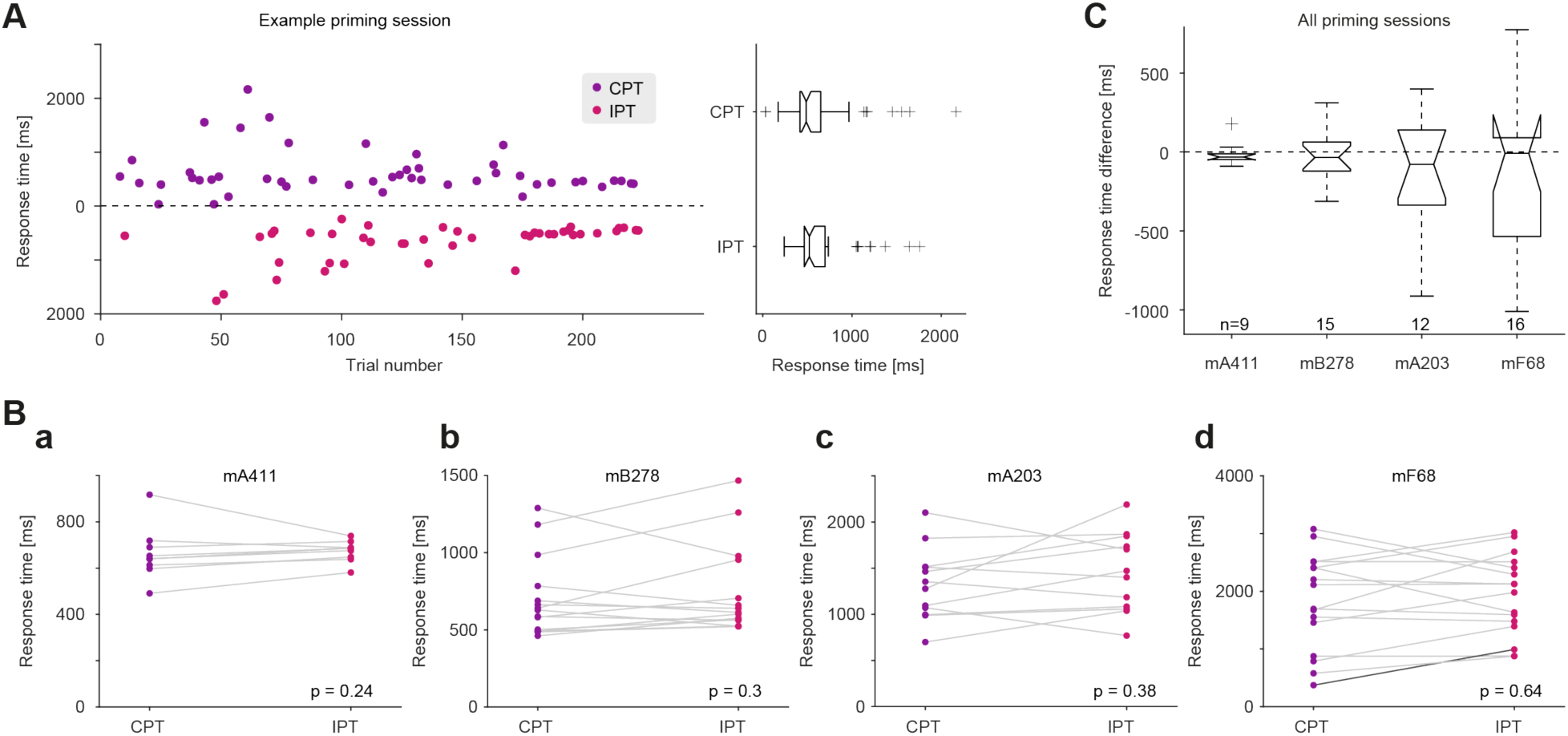
None of the mice show a priming effect in response times. **(A)** Response times during all PTs carried out in an example priming session performed by subject mB278 (same session as described in Fig. 5A). In a given PT, response time is defined as the duration from stimulus onset (target tone) to the response (side-port choice; Fig. 2C**, bottom**). **Left**, The response times in each trial of the session. The bottom half of the graph corresponds to the response times in IPTs (pink) and the top half of the graph corresponds to the response times in CPTs (violet). **Right**, Response times during the example session are not consistently different between n=48 CPTs and n=43 IPTs (p=0.15, Mann-Whitney U-test). **(B)** Median response times for all CPTs and IPTs during every priming session of every subject. P-values, two-tailed Wilcoxon’s signed-rank paired test. Sample sizes and all other conventions are the same as in Fig. 5C. **(C)** The differences of response times across all priming sessions for the four subjects. Box plot conventions are the same as in Fig. 3C.

At the session level, response times were not consistently longer or shorter during CPTs compared with IPTs during any session of subject mB278 (p>0.05 in all cases; n=15 sessions; light grey lines in **Fig. 6Bb**). At the subject level, there was no consistent difference between CPTs and IPTs in neither median response times (p=0.3, Wilcoxon’s paired signed-rank test) nor the variance of response times (p=0.79, permutation test; **Fig. 6Bb**). Similar results were observed for the other three mice (session level: light grey lines in **Fig. 6B**; subject level: p>0.05 in all cases; **Fig. 6B**). Of 52 priming sessions, we observed a consistent difference in response times in only one session (dark grey line in **Fig. 6Bd**). The global difference in response times was −35 ms [−285 76] ms (median [IQR] over n=52 priming sessions; **Fig. 6C**). Thus, none of the mice exhibited a priming effect in response times.

### The prime stimuli exert an effect on success rate via differential interference

The specificity of the priming effect to success rates leaves open the question of whether congruent prime stimuli facilitate performance, incongruent primes interfere with performance, or both. Since success rates during DTs were not perfect (median [IQR]: 88% [79% 91%]; n=84 discrimination sessions), there may be room for both improvement and deterioration. To contrast the possibilities, we compared performance in CPTs, IPTs, and DTs during the same session. In the example session of subject mB278, the animal made correct choices during 131/143 (92%) DTs, 47/48 (98%) CPTs, and 35/43 (81%) IPTs (**Fig. 7A**). The success rates during CPTs were higher than during DTs (p=0.005, Monte Carlo test; **Fig. 7A, right**), and success rates during IPTs were lower than during DTs (p=0.007). Thus, success rates during PTs may be either higher or lower than during same-session DTs.

**Figure 7.**
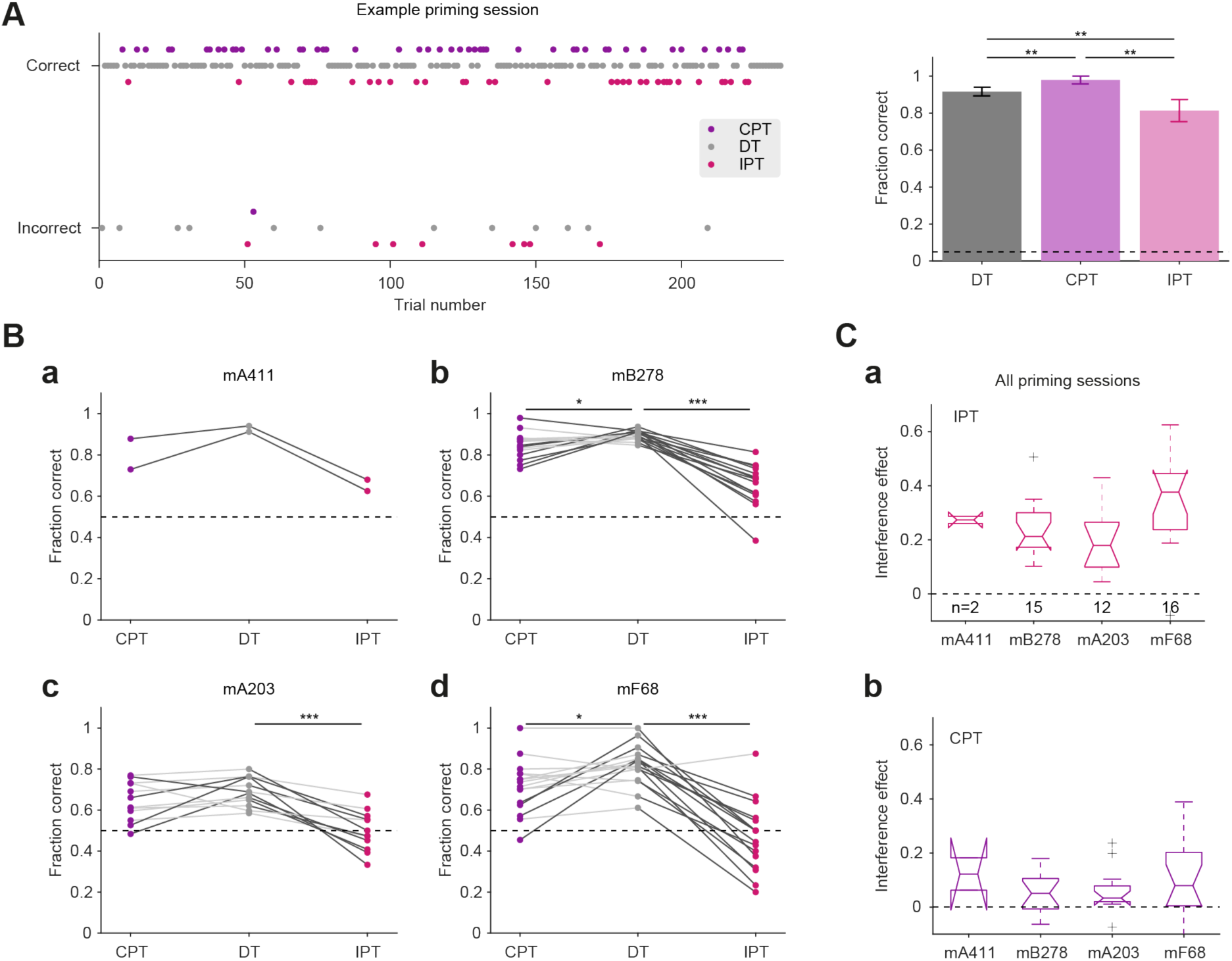
The prime stimuli exert an effect on success rate via differential interference. **(A)** All trials during an example priming session of mB278 (same session as described in Fig. 5A and Fig. 6A). CPTs, DTs, and IPTs are presented in a pseudorandom order. **Left**, Mouse choices. Trials with correct choices are shown on top, and trials with incorrect choices at the bottom. Stimulus frequency is not indicated. **Right**, Success rates during the three trial types. **: p<0.01, Monte Carlo test. **(B)** The success rates for all CPTs, DTs, and IPTs during every priming session of every subject. */**/***: p<0.05/p<0.01/p<0.001, two-tailed Wilcoxon’s signed-rank paired test. All other conventions are the same as in Fig. 5C. **(C)** The interference effect, defined as the session-specific difference between success rates during DTs and PTs. (**a**) Interference during IPTs. (**b**) Interference during CPTs.

To determine whether and in what direction performance is modified during CPTs and IPTs, we compared success rates at the subject level. For subject mB278, there was a consistent intra-session difference (p<0.05, Monte Carlo test) between IPT success rates compared to DT success rates during all 15/15 (100%) priming sessions (p<0.001, Wilcoxon’s paired test; dark grey lines at the right side of **Fig. 7Bb**). The median [IQR] interference was 21% [16% 31%] (n=15 sessions). In contrast, the difference between same-session CPTs and DTs was consistent during 7/15 (47%) of the sessions (p=0.03; dark grey lines at the left side of **Fig. 7Bb**) and the median [IQR] interference was 5.1% [-3.6% 11.6%]. Similar results were obtained for the other subjects (**Fig. 7B**). Thus, although both facilitation and interference are observed during priming trials, the dominant effect is interference.

Over all priming sessions for which data were available, the median [IQR] interference effect of the incongruent primes was 24% [17% 35%] (n=45 sessions in four mice; **Fig. 7Ca**). The incongruent primes modified success rates during 40/45 (89%) sessions. In all 40 sessions, the effect of IPTs was interference. The median effect of congruent primes was also interference, 5.4% [0.95% 12%] (**Fig. 7Cb**). However, in two sessions of two different subjects there was consistent facilitation (mB278, 6.3%; mA203, 7.4%). Thus, incongruent primes exert stronger interference compared with congruent primes, suggesting differential interference as a cognitive mechanism for short-term priming in mice.

## Discussion

We developed a short-term response priming paradigm for mice. Four subjects were trained on the auditory task, and all mice exhibited a priming effect in success rate. However, none of the mice exhibited a priming effect in response times. The prime stimuli exerted their effect via differential interference, predominantly during incongruent priming trials. The paradigm enables investigating the neurobiological basis of the short-term priming phenomenon with a new set of tools applicable to rodents.

### Comparison to human studies

In most human response priming paradigms, the priming effect is detected in response time^28,31,57,58^. Success rates remain at the ceiling, and are unmodified by the prime-target congruency. In other studies, the priming effect is found in both success rates and response times^36,38^. Then, the typical effect of the prime stimuli on success rates is reduction during IPTs, without modifying the already-perfect performance during CPTs^50^. Thus, response times are more sensitive than success rates in human response priming studies.

In the present mouse experiments, we found a priming effect only in success rates, providing two qualitative differences between human^36,38,50^ and mouse performance (**Fig. 1C**). First, in mice, the priming effect is in success rates and not in response times, whereas in humans, the priming effect is both in success rates and response times. Second, in mice, the main effect of the prime stimuli is interference, whereas in humans the main effect is interference following incongruent prime stimuli and improvement (or no effect) following congruent primes.

The first difference, namely the lack of an observed priming effect in mouse response times, may stem from the different task requirements or from the different setting and apparatus used for humans and mice. Physical requirements differ, since moving your entire body towards the correct port in every trial demands much more energy compared to pressing a button. Alternatively, it is possible that even when performing the exact same task with the exact same settings, humans and mice would have different time constants and exhibit different temporal variability. Finally, there could have been an effect in response times which was not detected.

The second difference, namely that the main effect of the prime stimuli on mouse performance is interference, may indicate that the mere presence of the additional prime stimulus is perceived as a distractor and hinders mouse behavior more than human behavior. Perceiving the prime as a distractor may stem from differences in some cognitive abilities between humans and mice. For instance, humans and mice may assign different attentional weights to different kinds of events, and may have different attitudes to the concept of a behavioral experiment.

### Extension to non-auditory and multi-modal priming

The current paradigm involves a specific auditory intra-modal short-term response priming task. However, the apparatus employed for the task was designed to be suitable for various extensions. First, the structure of the apparatus enables the development of a visual priming task in mice. For instance, blue light may be associated with left port choice, and green light with right port choice. Second, the apparatus also allows the potential study of inter-modal priming^29,58,59^, in which the prime stimulus would be auditory, and the target stimulus would be visual, or vice versa. Third, the apparatus allows extending the conscious priming paradigm to an unconscious priming paradigm^60–62^, for instance by using white noise to provide forward and/or backward masking of the prime stimuli.

### Extension to unconscious priming

Priming paradigms that include masks allow testing the influence of different types of unconscious processing on conscious performance^63–65^. Indeed, unconscious priming paradigms provide a window for researching unconscious processing pathways, that subjects cannot report. Because of the relative complexity of the paradigm, unconscious short-term memory is presently studied almost only in humans, limiting the questions that can be asked and mechanisms that can be studied. Thus, future extensions of the present mouse priming paradigm may allow studying unconscious processing in rodents.

A key issue during the study of unconscious processing is the necessity to verify that the stimulus was not perceived. With humans, the verification may involve a subjective report, but with animals, other methods are necessary. The mouse priming task can be extended to study unconscious processing as follows. Referring to **Fig. 2B**, trials during discrimination sessions (stage 1) can be supplemented with two consecutive auditory white noise masks before the target stimulus. During priming trials, the same white noise masks can be split into two stimuli that flank the prime. The validation task (stage 2), can be replaced with a calibration task in which the tones are preceded and followed by the same white noise masks. By reducing the duration of the tones and quantifying success rates, the longest ignored tone duration can be determined for every subject and utilized as the duration of the prime stimuli in the priming task (stage 3). An unconscious priming paradigm may enable the study of neural correlates of consciousness in freely-moving mice.

### Using the rodent paradigm to study the neurobiological basis of priming

Several types of memory, including implicit short-term memory, have not been studied with invasive techniques. Freely-moving mice have emerged as a robust model for studying various neuronal mechanisms using high density silicon probes recordings combined with optogenetics^66–68^. Here, we created a conscious short-term response priming paradigm for mice. We tested the paradigm and found a consistent priming effect of success rate in all subjects. We propose that the paradigm and the future extensions thereof provide a platform for researching the neurobiological mechanisms underlying short-term priming, using tools including genetic targeting, molecular analyses, and cellular-network electrophysiology. The present work forms a necessary first step for investigating the intracortical cellular-network level basis of short-term priming in particular, and of implicit short-term memory in general.

## Acknowledgements

We thank Liad Mudrik for useful discussions, and Samuel Frere, Liad Mudrik, Shimrit Oz, and Tali Remenick for constructive comments. This work was supported by European Research Council 679253; the Canadian Institutes of Health Research (CIHR), the International Development Research Centre (IDRC), the Israel Science Foundation (ISF), and the Azrieli Foundation 2558/18.

## Author contributions

S.S. and E.S. conceived and designed the experiments. S.S., H.E.S., and E.S. designed and constructed the experimental apparatus. S.S. carried out the experiments. S.S. and E.S. analyzed the data and wrote the manuscript with input from H.E.S.

## Materials and Methods

### Experimental animals

Four freely-moving adult mice of both sexes were used in this study (**Table S1**). The first three subjects were hybrid mice. Hybrid mice were used since compared to progenitors, hybrids exhibit reduced anxiety-like behavior and improved learning^67^. The first subject (mA411) was a hybrid female mouse, generated by crossing an FVB/NJ female (JAX #001800) and a C57BL/6J male (JAX #000664, The Jackson Laboratory). The second and third subjects, mB278 and mA203, were single-transgenic hybrid males, generated by crossing an FVB/NJ female with a parvalbumin (PV)-Cre male (JAX #008069). mB278 and mA203 were injected with a viral vector expressing Jaws^69^ (rAAV8/hSyn-Flex-Jaws) in neocortex and hippocampus as previously described^70^. The fourth subject, mF68, was a dual-transgenic male, generated by crossing a CaMKII-Cre female (JAX #005359) with an Ai32 male (JAX #012569). In the third and fourth mice, electrophysiological recordings and optical manipulations were carried out during some sessions. All animal handling procedures were in accordance with Directive 2010/63/EU of the European Parliament, complied with Israeli Animal Welfare Law (1994), and approved by the Tel Aviv University Institutional Animal Care and Use Committee (IACUC #01-16-051 and #01-21-012).

### Implanted probes

Two of the mice trained on the priming task were first implanted with a silicon diode-probe mounted on a micro-drive as previously described^70^. Subject mA203 was implanted with a single-shank, 32-site linear probe (20 μm inter-site spacing; A1×32-Edge, NeuroNexus) equipped with an optical fiber coupled to a red LD (638 nm). Subject mF68 was implanted with a dual-sided 6-shank/128-channel probe (15 μm inter-site spacing; Stark64 dual-sided, Diagnostic Biochips) equipped with two optical fibers coupled to blue LEDs (470 nm, on shanks 3 and 5). The diode-probes enable intra-cortical recordings during task performance, together with optical manipulations to silence (in mA203) or activate (in mF68) different types of neurons. The micro-drives allow changing probe location between sessions, enabling recordings from different depths along the vertical axis. Results of electrophysiological recordings are not included in the present report.

### Apparatus

All behavioral sessions were conducted with the subject inside an audio-visual behavioral apparatus (**Fig. 2A**). The apparatus consists of a 30 x 40 x 50 cm (L x W x H) open-top black-colored Plexiglas box. The front wall of the box hosts all sensors and actuators, arranged as three transparent behavioral ports. A speaker (22TAF/G, SEAS Prestige) for audio stimuli is located above the middle port. Every port contains a white LED for visual signaling; an infra-red LED-phototransistor pair for detecting nose pokes; and a steel tube connected via a flexible (Tygon) tube to a solenoid valve (003-0141-900, Parker) for water delivery. The solenoids were calibrated periodically to enable micro-liter resolution control of the amount of water dispensed.

The apparatus is controlled by a behavioral controller (BC). The BC is a metal enclosure containing a microcontroller (Arduino Mega) connected to custom-made electronic circuitry driven by a grounded power supply. Relevant digital events (sensor detection, actuator action, flow control) are routed from the BC via a multi-wire cable to the electrophysiology data acquisition system (DAQ) and to a real-time digital signal processor (DSP; RX8, Tucker-David Technologies). The microcontroller is connected to a PC via USB, and a custom graphical user interface is used to program the controller and report events to the experimenter. The direct communication between the BC, the DAQ, and the DSP allows precise temporal alignment of auditory stimuli, behavior, electrophysiological activity, and optogenetic stimuli.

### Water deprivation during training and testing

Throughout the behavioral training and testing period, the mice were kept under a water deprivation protocol during weekdays, and received free water during weekends. During weekdays, every mouse received at least 40 ml/kg of water (i.e., 1 ml/day for a 25 g mouse). Body weight was maintained above 85% at all times. During every behavioral session, the mouse was placed in the apparatus. The LED in the middle port was used to indicate trial onset to the mouse, and all three ports were used to quantify performance and govern the tasks in real time. At the beginning of the training period of the discrimination task (**Fig. 2B**), the reward volume was 10 μl for every successful trial, which was gradually decreased down to 6 μl. Subsequently, the water reward volume was the same during all tasks and at all ports.

### The discrimination task

The first stage of the learning and testing process (**Fig. 2B**) contained discrimination sessions. Every session contained two types of discrimination trials (DTs; **Fig. 2CD, top; video1**). The target stimuli were 300 ms pure tones with frequencies of 5 kHz (“low target tone”) or 10 kHz (“high target tone”). Correct performance involved the mouse poking a side port according to the tone played: the left port for the low target tone, and the right port for the high target tone. The stimulus-response contingency was the same for all mice. Before every trial, the LED of the middle port was on. To initiate a trial, the mouse had to poke the middle port to indicate readiness (**Fig. 2C, top**). 5 ms later, the target tone was played, the middle port LED turned off, the two side port LEDs turned on, and the system waited for the mouse to make a choice. If the mouse poked a side port or if 12000 ms elapsed without a side poke, the two side port LEDs turned off, the middle port LED turned on, and the system waited for the mouse to initiate the next trial. A correct choice of a side port resulted in water reward in that port. Lack of choice or an incorrect choice ended the trial without a water reward.

### The validation task

The second stage of the learning and testing process consisted of validation sessions (**Fig. 2B**). With human subjects, the irrelevance of the prime stimulus for correct performance of the task can be explained verbally but with animals, verbal explanations are not possible. To “explain” to the mouse the irrelevance of the priming stimulus for choosing a side port, we established a second learning stage termed “validation”. The purpose of the validation stage is to ensure that the mouse will understand that the short (prime) stimuli are not informative and should be ignored, and will keep responding to the long (target) stimuli.

The validation sessions consisted of a 3:1 pseudorandom mixture of DTs and validation trials (VTs; **Fig. 2CD, middle; video2**). The DTs were identical to the DTs employed during discrimination sessions, in which tone durations were 300 ms. In contrast, the tones in every VT were 15 ms pure tones with frequencies of 5 kHz (“short low tone”) or 10 kHz (“short high tone”). During a validation session, the mouse should ignore the short tones during the VTs, and respond to the long tones during the DTs exactly as during discrimination sessions. Ignoring the short tones can be done either by continued poking of the middle port until the next trial is initiated, or by leaving the middle port without selecting a side port. An alternative desirable response during VTs is random side port selection, i.e., a situation in which the not-ignored “success rate” is close enough to 0.5.

To initiate a VT the mouse had to poke the illuminated middle port, exactly as during a DT. Throughout the VT, the middle port LED was kept on and the two side ports LEDs were kept off. 5 ms after the VT was initiated, the short tone was played for 15 ms (**Fig. 2C, middle**) and the system waited for at least 1000 ms. If the mouse poked the middle port after the 1000 ms period has ended but before an additional 12000 ms have elapsed, a reward was given at the middle port. After a reward and a 1500 ms inter-trial interval, the mouse could initiate the next trial. Alternatively, if the mouse poked a side port at any point during the 13000 ms interval, the trial ended without a water reward. To prevent extinction of the discrimination task, a VT could only be followed by a DT (**Fig. 2D, middle**).

### The priming task

The third stage of the process contained priming sessions. Every priming session contained a 3:2 pseudorandom mixture of DTs and priming trials (PTs; **Fig. 2CD, bottom; video3**). The DTs were identical to the DTs employed during discrimination sessions. Every PT consisted of a pair of (prime, target) stimuli: the prime stimulus (short 15 ms tone, 5 kHz or 10 kHz) was followed by a target stimulus (300 ms tone, 5 kHz or 10 kHz; **Fig. 2C, bottom**). Thus, PTs could be either congruent (CPT; **Fig. 1B, left**) or incongruent (IPT; **Fig. 1B, right**). During a CPT, the stimulus pair had identical frequencies, {(5 kHz, 15 ms), (5 kHz, 300 ms)} or {(10 kHz, 15 ms), (10 kHz, 300 ms)} for left or right CPTs, respectively. During an IPT, the stimulus pair had different frequencies, {(10 kHz, 15 ms), (5 kHz, 300 ms)} or {(5 kHz, 15 ms), (10 kHz, 300 ms)} for left or right IPTs, respectively.

A PT began when the mouse poked the illuminated middle port. 5 ms later, the prime stimulus (a short 15 ms pure tone) was played. To prevent extinction of the validation task, a reward was given in the middle port at the same time. After a 200 ms delay, the target stimulus (a 300 ms pure tone) was played, the middle LED turned off, the two side port LEDs turned on, and the system waited up to 12000 ms for the mouse to make a choice. Thus, stimulus onset asynchrony (SOA) was fixed at 215 ms. If the mouse poked the correct side port, a water reward was given in the same port. When the mouse poked a side port, the side LEDs immediately turned off, the middle LED turned on, and the system waited for the mouse to initiate a new trial by a poking the middle port. The new trial could be a DT or a PT (**Fig. 2D, bottom**). If the mouse poked a side port before the target stimulus has been played, no reward was given, the LEDs reset, and the mouse could initiate a new trial.

### Real-time countering of automated strategies during the discrimination task

During learning of a new 2AFC task, mice tend to develop automated choice strategies which allow obtaining a reward in about 50% of the trials without actually learning the task^53,71^. Possible automated strategies include alternation (choosing the right and left ports alternately), perseveration (e.g., choosing the right port in every trial), win-stay/loose-shift, and win-shift/loose-stay. To prevent subjects from developing automated strategies during the discrimination task, we used online prediction of the automated strategy that the mouse is most likely to be taking. The algorithm^72^ estimated the instantaneous strategy and provided trials that minimize the effectiveness of the strategy. The procedure ensured that a mouse will not be able to quench its thirst without correctly performing the task.

### Quantification of performance during the discrimination task

We defined successful performance in a given discrimination session if (1) performance was above chance (Binomial test comparing to chance level of 0.5), and (2) the fraction of correct responses out of all (ignored and not-ignored) trials was above 0.85. We state that the mouse has finished the training phase and has learned the discrimination task (**Fig. 2B**) when at least four successful sessions were performed consecutively. Measurements of post-learning performance (testing sessions) were taken from the first of the four sessions and onwards.

### Quantification of performance during the validation task

To quantify performance in the validation task, we introduced an irrelevance score (**Fig. 4B**). The score quantifies the extent to which short tones are effectively ignored by the subject, and can be interpreted as how well the subject “understands” that short tones are irrelevant to the discrimination task. Typically, a mouse well-trained on the discrimination task that is exposed to the validation task for the first time, responds to the short tones as to long tones and pokes a side port immediately. During training on the validation task, the mouse is required to learn to ignore the short tones. The irrelevance score is defined as:

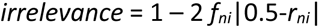

where *f_ni_* is the fraction of trials that were not ignored and *r_ni_* is the “success rate” during not-ignored trials. The irrelevance score is 1 if all short tones are ignored (*f_ni_*=0) or if “success rate” during all not-ignored trials is 0.5 (*r_ni_*=0.5; **Fig. 4B, red**). The irrelevance score is 0 if the mouse did not ignore any of the short tones (*f_ni_*=1) and has a not-ignored “success rate” of either *r_ni_*=0 or *r_ni_*=1 (**Fig. 4B, blue**).

In validation sessions, we defined chance level performance as a stimulus-dependent choice pattern during the VTs that was identical to the choice pattern during the DTs of that specific session. The rationale is that the default behavior of a mouse that has mastered the discrimination task and is exposed for the first time to a validation task is to treat short tones as target tones. We defined validation performance in a single session as successful if the irrelevance score was (1) above “chance” (Monte Carlo resampling test, comparing to chance level score based on the same-session DTs), and (2) above 0.85. The Monte Carlo resampling involved 10000 iterations, in which choices during randomized VTs were drawn from the empirical distribution of the stimulus-dependent choices during same-session DTs. We state that the mouse has finished the training phase and learned the validation task (**Fig. 2B**) when at least four successful validation sessions have been performed consecutively. Measurements of post-learning performance (testing sessions) were taken from the first of the four sessions and onwards.

### Quantification of performance during the priming task

The discrimination and validation tasks were designed to train animals on a specific stimulus-response association, and training continued until performance stabilized and specific criteria were achieved (**Fig. 2B**). In contrast, the priming task only involved testing the effect of short tones on the discrimination of long tones, and no performance criteria were set. A second difference is that the effect of the prime stimuli on discrimination during the priming task cannot be characterized at the single-trial level (correct/incorrect), but rather at the session level. We defined a “priming effect” in success rates as the difference between success rates during CPTs and success rates during same-session IPTs. To determine statistical significance, we used a permutation test, shuffling the labels of the CPTs and IPTs and comparing the observed priming effect (difference of success rates) with the null distribution of the priming effects obtained by 10000 iterations.

### Statistical analyses

In all statistical tests used in this study, a significance threshold of α=0.05 was used. All descriptive statistics (n, median, IQR, mean, SEM) can be found in the Results, figures, and figure legends. For DTs, we used a one-tailed Binomial test comparing to chance level, 0.5. For VTs, we used Monte Carlo randomization (10000 iterations), drawing choices from the empirical distribution of same-session DTs. For PTs, we used a permutation test (10000 iterations), shuffling the labels between same-session CPTs and IPTs. Differences between medians of two paired groups were tested using Wilcoxon’s signed-rank test (two tailed). Differences between medians of two unpaired groups were tested using Mann-Whitney’s U-test (two tailed). Differences between the variances of two groups were tested using a permutation test (two tailed). All statistical analyses were conducted in MATLAB (MathWorks).

## Tables

**Table S1.** Animal subjects.

| Animal ID | Strain<sup>1</sup> | Sex | Age<sup>2</sup> [weeks] | Weight<sup>2</sup> [g] | Probe |
| --- | --- | --- | --- | --- | --- |
| mA411 | HYB | Female | 8 | 19.4 |  |
| mB278 | HYB | Male | 12 | 30.5 |  |
| mA203 | HYB | Male | 32 | 32.8 | Linear32 |
| mF68 | C57 | Male | 27 | 29 | Stark64 dual-sided |

<sup>1</sup> HYB, hybrid mice, the F1 generation of an FVB/NJ female and a C57-derived male.
<sup>2</sup> At the beginning of the training period, excluding the weight of the implant.

## Video legends

### Video 1. Subject mF68 performing the discrimination task (stage 1)

In every trial, the pure tone stimulus is denoted as “Low DT” (5 kHz, 300 ms) or “High DT” (10 kHz, 300 ms). The response is denoted as “Correct”, “Incorrect”, or “Ignored”. For a 5 kHz / 10 kHz long tone, the correct response is poking the left / right port, respectively.

### Video 2. Subject mA203 performing the validation task (stage 2)

In every trial, the pure tone stimulus is denoted as “Low DT” (5 kHz, 300 ms), “High DT” (10 kHz, 300 ms), “Low VT” (5 kHz, 15 ms), or “High VT” (10 kHz, 15 ms). The response is denoted as “Correct”, “Incorrect”, or “Ignored”. For a 5 kHz / 10 kHz long tone (300 ms), the correct response is poking the left / right port, respectively. For short tones (15 ms), the correct response is poking the middle port (ignorance).

### Video 3. Subject mB278 performing the priming task (stage 3)

In every trial, the pure tone stimulus is denoted as “Low DT” (5 kHz, 300 ms), “High DT” (10 kHz, 300 ms), “Low CPT” {(5 kHz, 15 ms), (5 kHz, 300 ms)}, “High CPT” {(10 kHz, 15 ms), (10 kHz, 300 ms)}, “Low IPT” {(10 kHz, 15 ms), (5 kHz, 300 ms)}, or “High IPT” {(5 kHz, 15 ms), (10 kHz, 300 ms)}. The response is denoted as “Correct”, “Incorrect”, or “Ignored”. For a 5 kHz / 10 kHz long tone, the correct response is poking the left / right port, respectively.

## References

1. Yi, Y. (1990). The effects of contextual priming in print advertisements. J. Consum. Res. 17, 215–222. 10.1086/208551.

2. Bruce, V., Carson, D., Burton, A.M., and Kelly, S. (1998). Prime time advertisements: Repetition priming from faces seen on subject recruitment posters. Mem. Cogn. 26, 502–515. 10.3758/BF03201159.

3. Sofi, S.A., Nika, F.A., Shah, M.S., and Zarger, A.S. (2018). Impact of subliminal advertising on consumer buying behaviour: An empirical study on young Indian consumers. Glob. Bus. Rev. 19, 1580–1601. 10.1177/0972150918791378.

4. Ahmed (2020). Priming In marketing - 5 great examples to increase conversion rate. Digital Marketing Agency - ALLDGT. https://alldgt.com/priming-in-marketing/.

5. Dennis, A.R., Yuan, L. (IVY), Feng, X., Webb, E., and Hsieh, C.J. (2020). Digital nudging: Numeric and semantic priming in E-Commerce. J. Manag. Inf. Syst. 37, 39–65. 10.1080/07421222.2019.1705505.

6. Bargh, J.A. (2006). What have we been priming all these years? On the development, mechanisms, and ecology of nonconscious social behavior. Eur. J. Soc. Psychol. 36, 147–168. 10.1002/ejsp.336.

7. Bowden, E.M., Jung-Beeman, M., Fleck, J., and Kounios, J. (2005). New approaches to demystifying insight. Trends Cogn. Sci. 9, 322–328. 10.1016/j.tics.2005.05.012.

8. Martel, A., Arvaneh, M., Robertson, I., Smallwood, J., and Dockree, P. (2019). Distinct neural markers for intentional and unintentional task unrelated thought. bioRxiv, 705061. 10.1101/705061.

9. Korzeniewska, A., Wang, Y., Benz, H.L., Fifer, M.S., Collard, M., Milsap, G., Cervenka, M.C., Martin, A., Gotts, S.J., and Crone, N.E. (2020). Changes in human brain dynamics during behavioral priming and repetition suppression. Prog. Neurobiol. 189, 101788. 10.1016/j.pneurobio.2020.101788.

10. Maftei, A., and Holman, A.C. (2022). Moral in the future, better now: Moral licensing versus behavioral priming in children and the moderating role of psychological distance. Curr. Psychol. 10.1007/s12144-022-03063-5.

11. Kirmani, A., Lee, M.P., and Yoon, C. (2004). Procedural priming effects on spontaneous inference formation. J. Econ. Psychol. 25, 859–875. 10.1016/j.joep.2003.09.003

12. Lempert, K.M., and Phelps, E.A. (2014). Chapter 12 - Neuroeconomics of emotion and decision making. In Neuroeconomics (Second Edition), P. W. Glimcher and E. Fehr, eds. (Academic Press), pp. 219–236. 10.1016/B978-0-12-416008-8.00012-7.

13. Liang, X., Xiao, F., Lei, Y., Li, H., and Chen, Q. (2020). N400/frontal negativity reveals the controlled processes of taxonomic and thematic relationships in semantic priming for artifacts. Psychophysiology 57, e13486. 10.1111/psyp.13486.

14. Over, H., and Carpenter, M. (2009). Eighteen-month-old infants show increased helping following priming with affiliation. Psychol. Sci. 20, 1189–1193. 10.1111/j.1467-9280.2009.02419.x.

15. Ahnert, L., Milatz, A., Kappler, G., Schneiderwind, J., and Fischer, R. (2013). The impact of teacher– child relationships on child cognitive performance as explored by a priming paradigm. Dev. Psychol. 49, 554–567. 10.1037/a0031283.

16. Campbell, M.C., Manning, K.C., Leonard, B., and Manning, H.M. (2016). Kids, cartoons, and cookies: Stereotype priming effects on children’s food consumption. J. Consum. Psychol. 26, 257–264. 10.1016/j.jcps.2015.06.003.

17. Sekerina, I.A., Fernández, E.M., and Clahsen, H. (2008). Developmental Psycholinguistics: On-line Methods in Children’s Language Processing (John Benjamins Publishing). 10.1075/lald.44.

18. Mandera, P., Keuleers, E., and Brysbaert, M. (2017). Explaining human performance in psycholinguistic tasks with models of semantic similarity based on prediction and counting: A review and empirical validation. J. Mem. Lang. 92, 57–78. 10.1016/j.jml.2016.04.001.

19. Forgas, J.P., Laham, S.M., and Vargas, P.T. (2005). Mood effects on eyewitness memory: Affective influences on susceptibility to misinformation. J. Exp. Soc. Psychol. 41, 574–588. 10.1016/j.jesp.2004.11.005.

20. Koegel, L.K., Koegel, R.L., Frea, W., and Green-Hopkins, I. (2003). Priming as a method of coordinating educational services for students with autism. LSHSS 34, 228–235. 10.1044/0161-1461(2003/019).

21. Lamb, R., Akmal, T., and Petrie, K. (2015). Development of a cognition-priming model describing learning in a STEM classroom. J. Res. Sci. Teach. 52, 410–437. 10.1002/tea.21200.

22. Mele, A.R. (2014). Surrounding free will: Philosophy, psychology, neuroscience (Oxford University Press). 10.1093/acprof:oso/9780199333950.001.0001.

23. Romero, F. (2016). Can the behavioral sciences self-correct? A social epistemic study. Stud. Hist. Philos. Sci. A. 60, 55–69. 10.1016/j.shpsa.2016.10.002.

24. Tsalikis, J. (2015). The effects of priming on business ethical perceptions: A comparison between two cultures. J. Bus. Ethics 131, 567–575. 10.1007/s10551-014-2243-3.

25. Was, C., Woltz, D., and Hirsch, D. (2019). Memory processes underlying long-term semantic priming. Mem. Cogn. 47, 313–325. 10.3758/s13421-018-0867-8.

26. Beer, A.L., and Diehl, V.A. (2001). The role of short-term memory in semantic priming. J. Gen. Psychol. 128, 329–350. 10.1080/00221300109598915.

27. Meyer, D.E., and Schvaneveldt, R.W. (1971). Facilitation in recognizing pairs of words: Evidence of a dependence between retrieval operations. J. Exp. Psychol. 90, 227–234. 10.1037/h0031564.

28. Schacter, D.L., and Buckner, R.L. (1998). Priming and the brain. Neuron 20, 185–195. 10.1016/S0896-6273(00)80448-1.

29. Graf, P., Shimamura, A.P., and Squire, L.R. (1985). Priming across modalities and priming across category levels: Extending the domain of preserved function in amnesia. J. Exp. Psychol. Learn. Mem. Cogn. 11, 386–396. 10.1037//0278-7393.11.2.386.

30. Buchner, A., Zabal, A., and Mayr, S. (2003). Auditory, visual, and cross-modal negative priming. Psychon. Bull. Rev. 10, 917–923. 10.3758/BF03196552.

31. Dehaene, S., and Changeux, J.P. (2011). Experimental and theoretical approaches to conscious processing. Neuron 70, 200–227. 10.1016/j.neuron.2011.03.018.

32. Dehaene, S., Naccache, L., Clec’H, G.L., Koechlin, E., Mueller, M., Dehaene-Lambertz, G. van de Moortele, P.F., and Le Bihan, D. (1998). Imaging unconscious semantic priming. Nature 395, 597–600. 10.1038/26967.

33. Kouider, S., and Dehaene, S. (2007). Levels of processing during non-conscious perception: A critical review of visual masking. Philos. Trans. R. Soc. Lond. B Biol. Sci. 362, 857–875. 10.1098/rstb.2007.2093.

34. Finkbeiner, M. (2011). Subliminal priming with nearly perfect performance in the prime-classification task. Atten. Percept. Psychophys. 73, 1255–1265. 10.3758/s13414-011-0088-8.

35. Rosenbaum, D.A., and Kornblum, S. (1982). A priming method for investigating the selection of motor responses. Acta Psychologica 51, 223–243. 10.1016/0001-6918(82)90036-1.

36. Schmidt, F., Haberkamp, A., and Schmidt, T. (2011). Dos and don’ts in response priming research. Adv. Cogn. Psychol. 7, 120–131. 10.2478/v10053-008-0092-2.

37. Chen, Y.C., and Spence, C. (2018). Dissociating the time courses of the cross-modal semantic priming effects elicited by naturalistic sounds and spoken words. Psychon. Bull. Rev. 25, 1138–1146. 10.3758/s13423-017-1324-6.

38. Schmidt, T., and Schmidt, F. (2009). Processing of natural images is feedforward: A simple behavioral test. Atten. Percept. Psychophys. 71, 594–606. 10.3758/APP.71.3.594.

39. Blough, P.M. (1989). Attentional priming and visual search in pigeons. J. Exp. Psychol. Anim. Behav. Process. 15, 358–365. 10.1037/0097-7403.15.4.358.

40. Brodbeck, D.R. (1997). Picture fragment completion: Priming in the pigeon. J. Exp. Psychol. Anim. Behav. Process. 23, 461–468. 10.1037/0097-7403.23.4.461.

41. Fremouw, T., Herbranson, W.T., and Shimp, C.P. (1998). Priming of attention to local or global levels of visual analysis. J. Exp. Psychol. Anim. Behav. Process. 24, 278–290. 10.1037/0097-7403.24.3.278.

42. Blough, D.S. (2000). Effects of priming, discriminability, and reinforcement on reaction-time components of pigeon visual search. J. Exp. Psychol. Anim. Behav. Process. 26, 50–63. 10.1037/0097-7403.26.1.50.

43. Nieder, A., Wagener, L., and Rinnert, P. (2020). A neural correlate of sensory consciousness in a corvid bird. Science 369, 1626–1629. 10.1126/science.abb1447.

44. Amitai, N., Weber, M., Swerdlow, N.R., Sharp, R.F., Breier, M.R., Halberstadtr, A.L., and Young, J.W. (2013). A novel visuospatial priming task for rats with relevance to Tourette syndrome and modulation of dopamine levels. Neurosci. Biobehav. Rev. 37, 1139–1149. 10.1016/j.neubiorev.2012.09.007.

45. Amitai, N., Powell, S., Weber, M., Swerdlow, N.R., and Young, J.W. (2015). Negative visuospatial priming in isolation-reared rats: Evidence of resistance to the disruptive effects of amphetamine. Cogn. Affect. Behav. Neurosci. 15, 901–914. 10.3758/s13415-015-0369-0.

46. Ruiz, A., Gómez, J.C., Roeder, J.J., and Byrne, R.W. (2009). Gaze following and gaze priming in lemurs. Anim. Cogn. 12, 427–434. 10.1007/s10071-008-0202-z.

47. McMahon, D.B.T., and Olson, C.R. (2007). Repetition suppression in monkey inferotemporal cortex: Relation to behavioral priming. J. Neurophysiol. 97, 3532–3543. 10.1152/jn.01042.2006.

48. Ben-Haim, M.S., Dal Monte, O., Fagan, N.A., Dunham, Y., Hassin, R.R., Chang, S.W.C., and Santos, L.R. (2021). Disentangling perceptual awareness from nonconscious processing in rhesus monkeys (Macaca mulatta). Proc. Natl. Acad. Sci. USA 118, e2017543118. 10.1073/pnas.2017543118.

49. Dehaene, S., Changeux, J.-P., and Naccache, L. (2011). The global neuronal workspace model of conscious access: From neuronal architectures to clinical applications. In Characterizing consciousness: From cognition to the clinic? Research and Perspectives in Neurosciences., S. Dehaene and Y. Christen, eds. (Springer), pp. 55–84. 10.1007/978-3-642-18015-6_4.

50. Vorberg, D., Mattler, U., Heinecke, A., Schmidt, T., and Schwarzbach, J. (2003). Different time courses for visual perception and action priming. Proc. Natl. Acad. Sci. USA 100, 6275–6280. 10.1073/pnas.0931489100.

51. Whyte, C.J., and Smith, R. (2021). The predictive global neuronal workspace: A formal active inference model of visual consciousness. Prog. Neurobiol. 199, 101918. 10.1016/j.pneurobio.2020.101918

52. Kiefer, M., Adams, S.C., and Zovko, M. (2012). Attentional sensitization of unconscious visual processing: Top-down influences on masked priming. Adv. Cogn. Psychol. 8, 50–61. 10.5709/acp-0102-4.

53. Jaramillo, S., and Zador, A.M. (2014). Mice and rats achieve similar levels of performance in an adaptive decision-making task. Front. Syst. Neurosci. 8, 173. 10.3389/fnsys.2014.00173.

54. Turner, J.G., Parrish, J.L., Hughes, L.F., Toth, L.A., and Caspary, D.M. (2005). Hearing in laboratory animals: Strain differences and nonauditory effects of noise. Comp. Med. 55, 12–23.

55. Pickering, M.J., and Ferreira, V.S. (2008). Structural priming: A critical review. Psychol. Bull. 134, 427–459. 10.1037/0033-2909.134.3.427.

56. Janiszewski, C., and Wyer, R.S. (2014). Content and process priming: A review. J. Consum. Psychol. 24, 96–118. 10.1016/j.jcps.2013.05.006.

57. Tulving, E., and Schacter, D.L. (1990). Priming and human memory systems. Science 247, 301–306. 10.1126/science.2296719.

58. Schacter, D.L. (1992). Priming and multiple memory systems: Perceptual mechanisms of implicit memory. J. Cogn. Neurosci. 4, 244–256. 10.1162/jocn.1992.4.3.244.

59. Kirsner, K., Milech, D., and Stumpfel, V. (1986). Word and picture identification: Is representational parsimony possible? Mem. Cognit. 14, 398–408. 10.3758/BF03197015.

60. Babiloni, C., Vecchio, F., Rossi, S., De Capua, A., Bartalini, S., Ulivelli, M., and Rossini, P.M. (2007). Human ventral parietal cortex plays a functional role on visuospatial attention and primary consciousness. A repetitive transcranial magnetic stimulation study. Cereb. Cortex. 17, 1486–1492. 10.1093/cercor/bhl060.

61. King, J.A., Korb, F.M., and Egner, T. (2012). Priming of control: Implicit contextual cuing of top-down attentional set. J. Neurosci. 32, 8192–8200. 10.1523/JNEUROSCI.0934-12.2012.

62. Keane, M.M., Cruz, M.E., and Verfaellie, M. (2015). Attention and implicit memory: Priming-induced benefits and costs have distinct attentional requirements. Mem. Cognit. 43, 216–225. 10.3758/s13421-014-0464-4.

63. Greenwald, A.G., Draine, S.C., and Abrams, R.L. (1996). Three cognitive markers of unconscious semantic activation. Science 273, 1699–1702. 10.1126/science.273.5282.1699.

64. Breitmeyer, B.G. (2015). Psychophysical “blinding” methods reveal a functional hierarchy of unconscious visual processing. Conscious. Cogn. 35, 234–250. 10.1016/j.concog.2015.01.012.

65. Mudrik, L., and Deouell, L.Y. (2022). Neuroscientific evidence for processing without awareness. Annu. Rev. Neurosci. 45, 403–423. 10.1146/annurev-neuro-110920-033151.

66. Stark, E., Roux, L., Eichler, R., and Buzsáki, G. (2015). Local generation of multineuronal spike sequences in the hippocampal CA1 region. Proc. Natl. Acad. Sci. USA 112, 10521–10526. 10.1073/pnas.1508785112.

67. Sloin, H.E., Bikovski, L., Levi, A., Amber-Vitos, O., Katz, T., Spivak, L., Someck, S., Gattegno, R., Sivroni, S., Sjulson, L., et al. (2022). Hybrid offspring of C57BL/6J mice exhibit improved properties for neurobehavioral research. eNeuro 9, ENEURO.0221-22.2022. 10.1523/ENEURO.0221-22.2022.

68. Levi, A., Spivak, L., Sloin, H.E., Someck, S., and Stark E. (2022). Error correction and improved precision of spike timing in converging cortical networks. Cell. Rep. 40, 111383. 10.1016/j.celrep.2022.111383.

69. Chuong, A.S., Miri, M.L., Busskamp, V., Matthews, G.A.C., Acker, L.C., Sørensen, A.T., Young, A., Klapoetke, N.C., Henninger, M.A., Kodandaramaiah, S.B., et al. (2014). Noninvasive optical inhibition with a red-shifted microbial rhodopsin. Nat. Neurosci. 17, 1123–1129. 10.1038/nn.3752.

70. Noked, O., Levi, A., Someck, S., Amber-Vitos, O., and Stark, E. (2021). Bidirectional optogenetic control of inhibitory neurons in freely-moving mice. IEEE Trans. Biomed. Eng. 68, 416–427. 10.1109/TBME.2020.3001242.

71. Carandini, M., and Churchland, A.K. (2013). Probing perceptual decisions in rodents. Nat. Neurosci. 16, 824–831. 10.1038/nn.3410.

72. Stoneking, C.J. (2018). Antibias; https://github.com/cjstoneking/antibias.

